# A novel fibrinogen gamma-chain mutation, p. γAla327Val, causes structural abnormality of D region and ultimately leads to congenital dysfibrinogenemia

**DOI:** 10.1101/2020.01.29.925172

**Authors:** Aiqiu Wei, Yangyang Wu, Liqun Xiang, Jie Yan, Peng Cheng, Donghong Deng, Faquan Lin

## Abstract

Congenital dysfibrinogenemia (CD) is a coagulation disorder caused by mutations in the fibrinogen gene, which result in abnormal fibrinogen function. Many studies have confirmed that over half of dysfibrinogenemia cases are asymptomatic. In this study, we aimed to investigate the pathogenesis of CD caused by γ Ala327Val heterozygous mutation, a new mutation, by studying fibrinogen function. Blood samples of patients were collected and the coagulation function, fibrinogen (Fg) aggregation test, fibrin clot lysis test, and SDS-PAGE were performed. Coagulation was monitored using a thromboelastometer, and the fibrin clot network structure was observed by scanning electron microscopy. The effect of the mutation on fibrinogen structure and function was predicted by molecular modeling. The fibrinogen activity concentration in patients with CD was significantly lower than that in healthy individuals. Thromboelastography showed that the K value of patients with CD was higher than that for healthy individuals. The Angle values were also decreased. The function of fibrinogen in patients with CD was low. Compared to fibrinogen from healthy individuals, fibrin size was different, the fiber network structure was loose, the pore size was increased, and the fiber branch nodes were increased for fibrinogen isolated from the proband. The γ Ala327Val mutation led to changes in the structure of fibrinogen D region, affecting its structural stability. Ala327Val heterozygous missense mutation in exon 8 of FGG gene γ-chain thus leads to abnormal fibrinogen structure and impairs the aggregation function of fibrinogen. This mutation is reported here for the first time.

## Introduction

Fibrinogen (Fg), also known as coagulation factor I, is mainly produced by liver cells and is secreted into peripheral blood. Its plasma concentration is 2-4 g/L [1]. Fibrinogen is involved in the formation of fibrin clots and platelet aggregation, and plays an important role in hemostasis. At the initial stage of fibrin clot formation, fibrinogen binds to platelet membrane protein IIb/IIIa, activates platelets, and mediates platelet aggregation, thus participating in the coagulation process. The relative molecular weight of fibrinogen is 340 kD; it is composed of two identical subunits, each of which comprises three different polypeptide chains (A α, B β, and γ); these two subunits are linked by disulfide bonds [2]. Fibrinogen molecules have three major functional regions, the central region (E region) and two symmetric distal spherical D regions (Figure 1). The three polypeptide chains A α, B β, and γ are encoded by FGA, FGB, and FGG genes (on chromosome 4), respectively [3].

**Figure 1.**
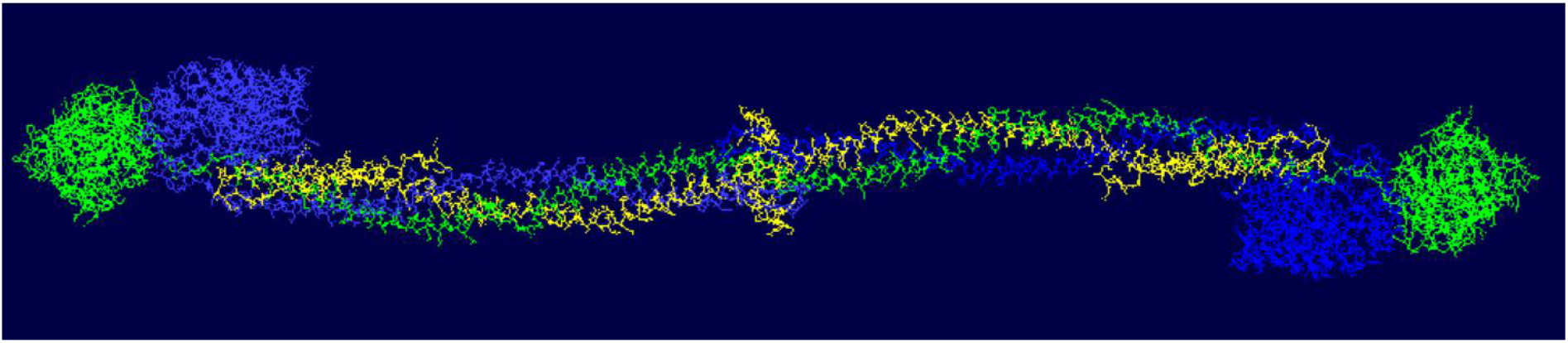
Molecular structure of fibrinogen.

Mutations in FGA, FGB, or FGG may lead to the development of congenital dysfibrinogenemia (CD). CD is a congenital blood disease caused by defects in fibrinogen genes, which lead to abnormal structure and function of fibrinogen molecules, and may affect coagulation. The clinical manifestations of CD are diverse; most patients with CD are asymptomatic [4], but a small number of patients with CD have thrombosis [5, 6] and bleeding events, and some patients have pulmonary hypertension and other symptoms [7]. As most patients with CD are asymptomatic, and the single detection method used currently in clinical laboratories may lead to misdiagnosis or missed diagnosis, the incidence of CD is thus difficult to determine at present.

In this study, we identified a novel heterozygous mutation (FGG c.1058C > T, γAla327Val) in an asymptomatic patient with abnormal fibrinogenemia. Functional studies of fibrinogen isolated from the patient and her family members were performed to study the molecular pathogenesis of CD caused by γ Ala327Val heterozygous missense mutation.

## Methods

### Basic data of patients and routine examination of coagulation function

The proband was a 45-year-old female patient. When she came to the hospital for physical examination, her coagulation function test result was abnormal: fibrinogen activity concentration was 0.75 g/L (Clauss method) and antigen concentration (immune turbidimetry method) was 1.59 g/L (normal reference range for both parameters: 2.0-4.0 g/L). Fibrinogen activity concentration was significantly lower than fibrinogen antigen concentration. Liver function, kidney function, and blood routine tests were normal, and there were no bleeding or thrombotic events in daily life. Four members of her family were examined for coagulation function and other parameters.

DNA was extracted from the peripheral blood samples of the proband and four members of her family. DNA sequencing was performed by Beijing Liuhe Huada Gene Technology Co., Ltd, Beijing, China.

### Fibrinogen aggregation test

Venous blood (5 ml, in sodium citrate anticoagulant) was drawn from the patients and healthy controls, and centrifuged at 4°C, 3000 rpm, for 15 min to separate the plasma. Then, 1 ml sodium citrate (0.09 mol/L) and 666 μl saturated ammonium sulfate (pH 5.5) were added to 1 ml plasma, and the mixture was incubated at 25°C for 70 minutes. After centrifugation, the supernatant was discarded and fibrinogen was obtained. The reaction system of the fibrinogen aggregation test was as follows: Fibrinogen 0.5 mg/ml (final concentration), 5 ml of 2 mol/L NaCl, 5 ml of 2 mmol/L CaCl2, and 20 mmol/L HEPES buffer to make the volume up to 90 ml. Then, 10 U/ml of 10 μl thrombin was added and the optical density (OD) of the sample at 365 nm was continuously monitored (one reading every 20 seconds for 30 minutes) using a multimode plate reader Multiskan Go (ThermoFischer). The aggregation curve was drawn according to the time and the corresponding OD value.

### Fibrin clot dissolution test

Fibrin clot dissolution test was performed by adding fibrinolytic enzyme and tPA into the fibrinogen aggregation test system to activate fibrin clot dissolution, and we continuously measured the OD value of the reaction system to reflect the rate of fibrin clot dissolution. The reaction system used for this test was as follows: The final concentrations of fibrinogen, thrombin, plasminogen, tPA, CaCl2, and NaCl were 0.5 mg/ml, 0.5 U/ml, 0.12 U/ml, 0.1 mg/ml, 8 mmol/L, 0.12 mmol/L, and 20 mmol/L, respectively. Finally, 20 mmol/L HEPES was used to make up the volume of the reaction system to 200 ml. The OD values of the samples at 365 nm were continuously monitored with Multiskan Go (ThermoFisher, USA) for 30 minutes at 20 s intervals. The fibrin clot dissolution curve was drawn according to the time and the corresponding OD value.

### Thromboelastography

Whole blood samples (1 ml) from the study subjects were added to the reagent bottle (Shaanxi Yuze Yi Medical Technology Co., Ltd Shaanxi, China); Then, 340 μl of the sample with 20 μl of 0.2 mol/L CaCl2 was added into the detection cup of the thromboelastography (TEG) instrument (Shaanxi Yuze Yi Medical Technology Co., Ltd, Shaanxi, China). As fibrinogen began to aggregate, the probe placed in the detection cup was subjected to shear stress produced during the formation and dissolution of the blood clot. The generated signal was transmitted to the processor in the form of a current and a TEG curve was obtained.

### SDS-PAGE

Fibrinogen was mixed with loading buffer (containing β-mercaptoethanol) and the mixture was heated for 5 min at 100°C. Sodium dodecyl sulfate polyacrylamide gel electrophoresis (SDS-PAGE) was performed on 10% gels using the BioRad Mini-Protean Tetra Electrophoresis System (Bio-Rad, CA, USA). The gel was stained with Coomassie brilliant blue R-250 and was scanned on a gel imager (Bio-Rad Laboratories (Shanghai) Co., Ltd.).

### Scanning electron microscopy

The proband and healthy individuals were administered 33 μl of fibrinogen each, and thrombin was added at a final concentration of 2 U/ml). After incubation for 3 hours at 37°C, the samples from proband and healthy individuals were rinsed with PBS buffer (pH 7.4, 0.1 mol/L), and incubated with 3% glutaraldehyde for 2 hours. Glutaraldehyde was discarded, samples were rinsed with PBS solution, and dehydrated with alcohol. The ultrastructure of the fibrin clot was observed with VEGA3LMU scanning electron microscope (TESCAN, Czech Republic).

### Modeling and analysis of amino acid mutation

The amino acid sequence of fibrinogen was obtained from the NCBI database, and a model of the fibrinogen structure was constructed using SWISS-MODEL website (https://swissmodel.expasy.org/)/. Swiss-Pdb Viewer software was used to analyze the effect of γ Ala327Val mutation on the function of fibrinogen.

## Results

### Routine coagulation function tests

The Results of the laboratory examination for the proband including electrolyte, liver and kidney function, and blood routine tests were normal. Examination of these parameters for other members of the family showed that the coagulation results of the proband’s sister, two brothers, and proband’s daughter were similar to those of the proband, and there were no abnormalities in the results of the electrolyte, liver and kidney function, and blood routine tests. There were no clinical manifestations in daily life. The results of the coagulation function test are given in table 1. A single base heterozygous missense mutation (C. 1058C > T, γ Ala327Val) in exon 8 of the fibrinogen gene was found in the proband and her family members (Figure 2).

**Figure 2.**
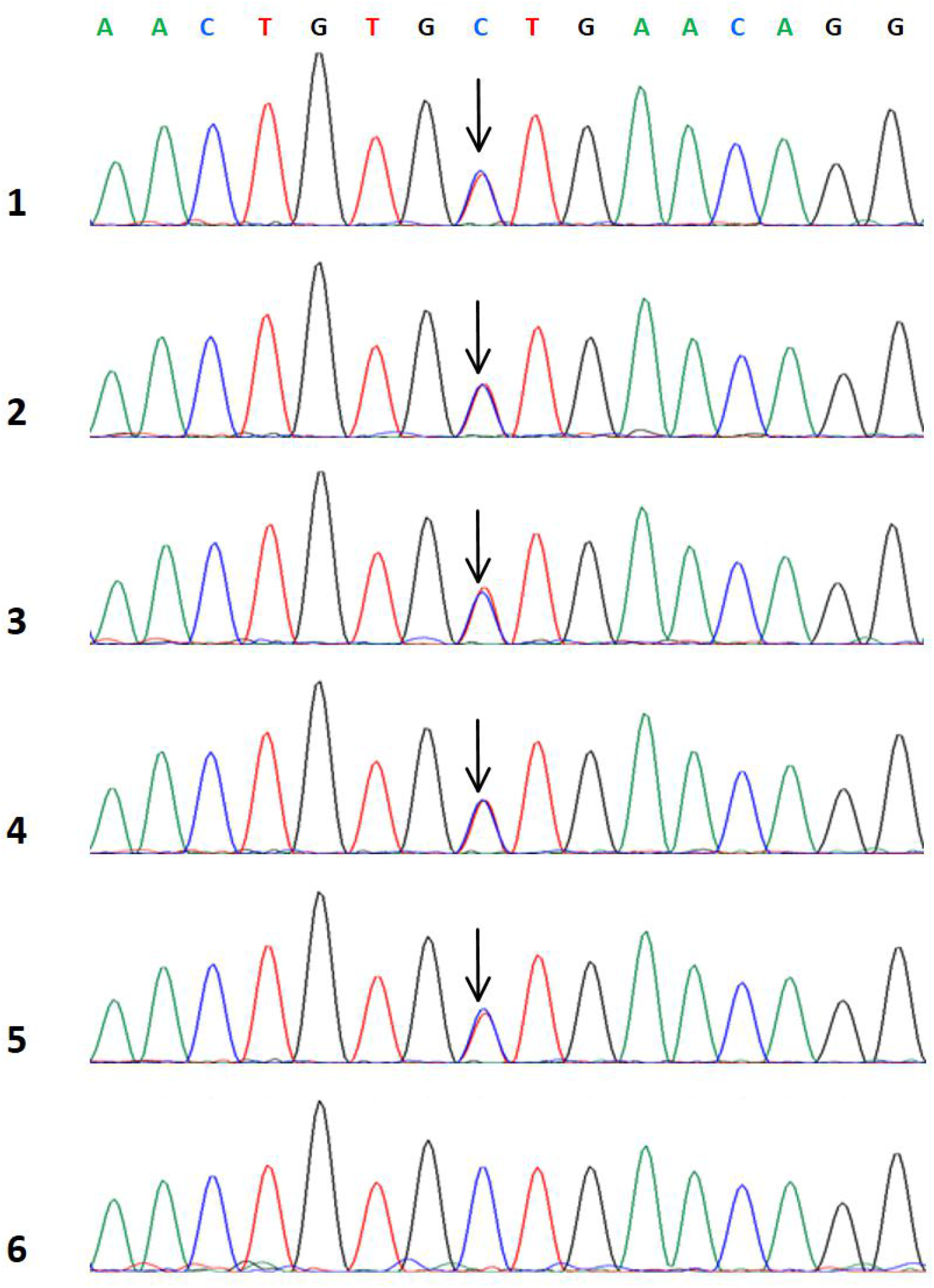
Sequences of gene.1: Proband; 2-5: Proband’s family member; 6: Healthy individual (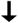: FGG gene c.1058C>T heterozygous mutation).

**Table 1.**
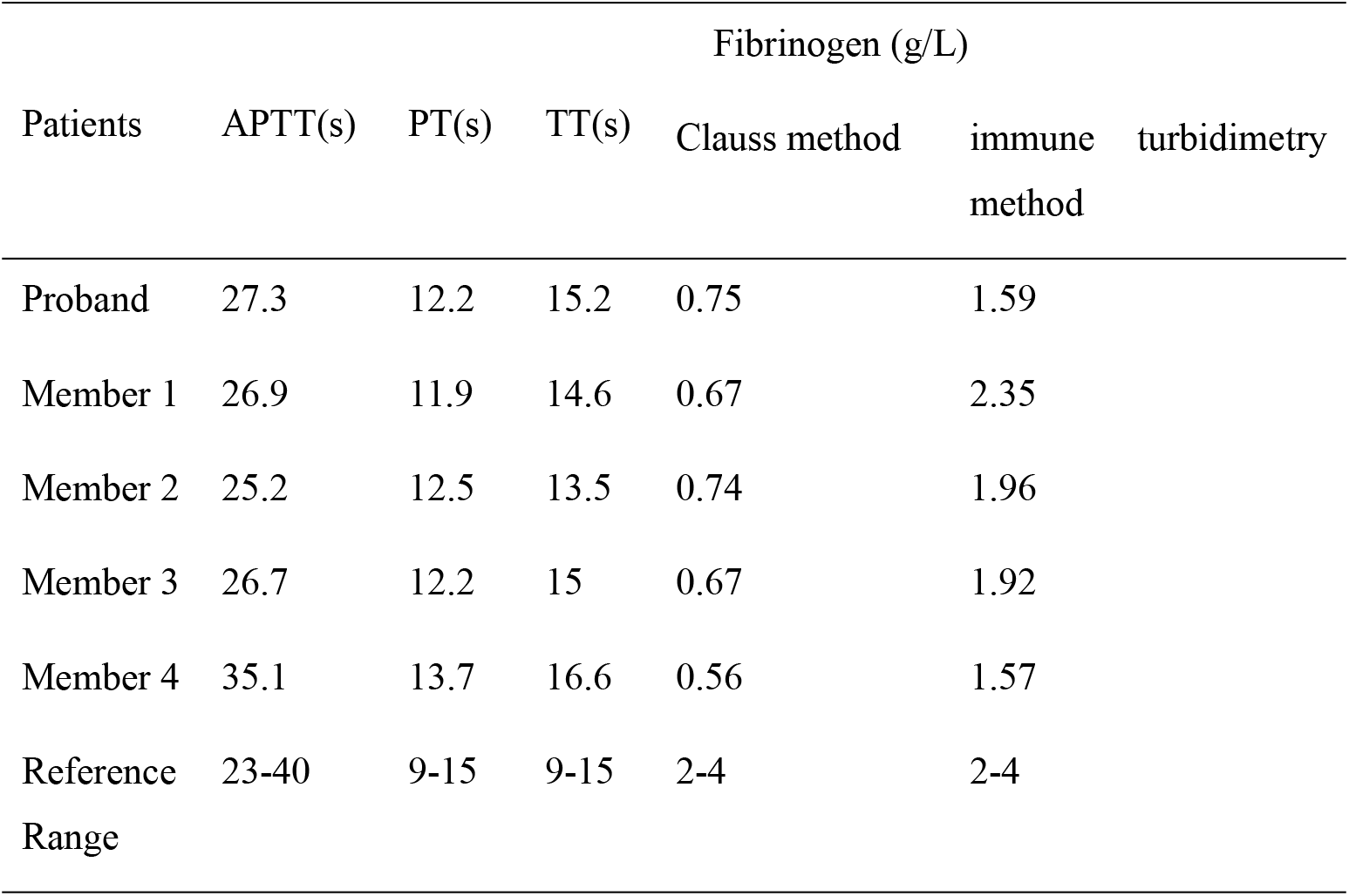
Coagulation function of proband and her family members.

### Fibrinogen aggregation test

Compared with the healthy individuals, the OD values of fibrinogen aggregation curves for fibrinogen isolated from the proband and her family members were changed slightly, and the maximum OD values from 5 patients (average 0.283) were lower than those of the healthy individual (0.422) (Figure 3).

**Figure 3.**
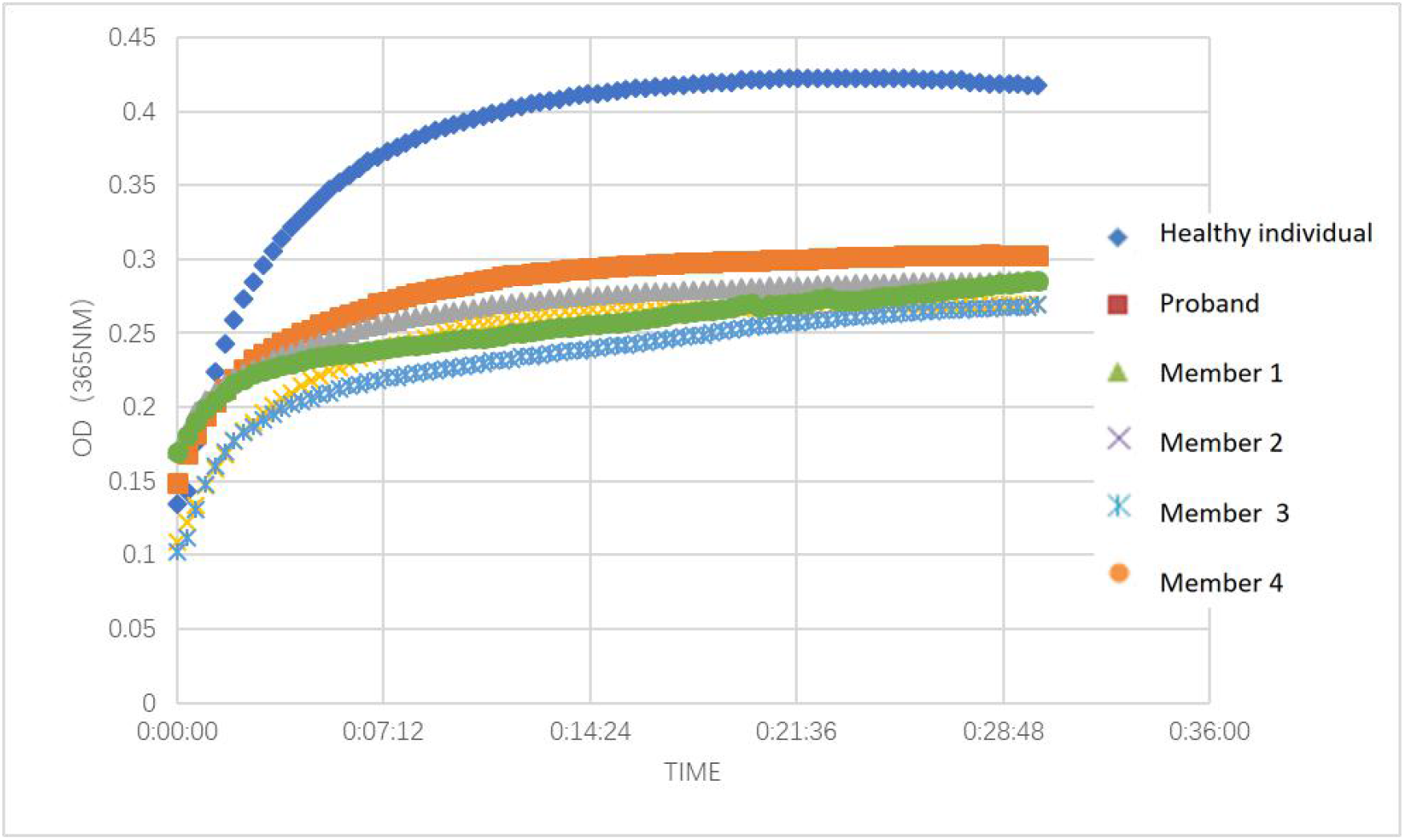
Fibrin polymerization curves.

### Fibrin clot dissolution test

The clot turbidity began to decrease after 496 seconds on average (healthy individuals: after 660 seconds). There was no fibrinolysis delay and resistance during clot dissolution (Figure 4).

**Figure 4.**
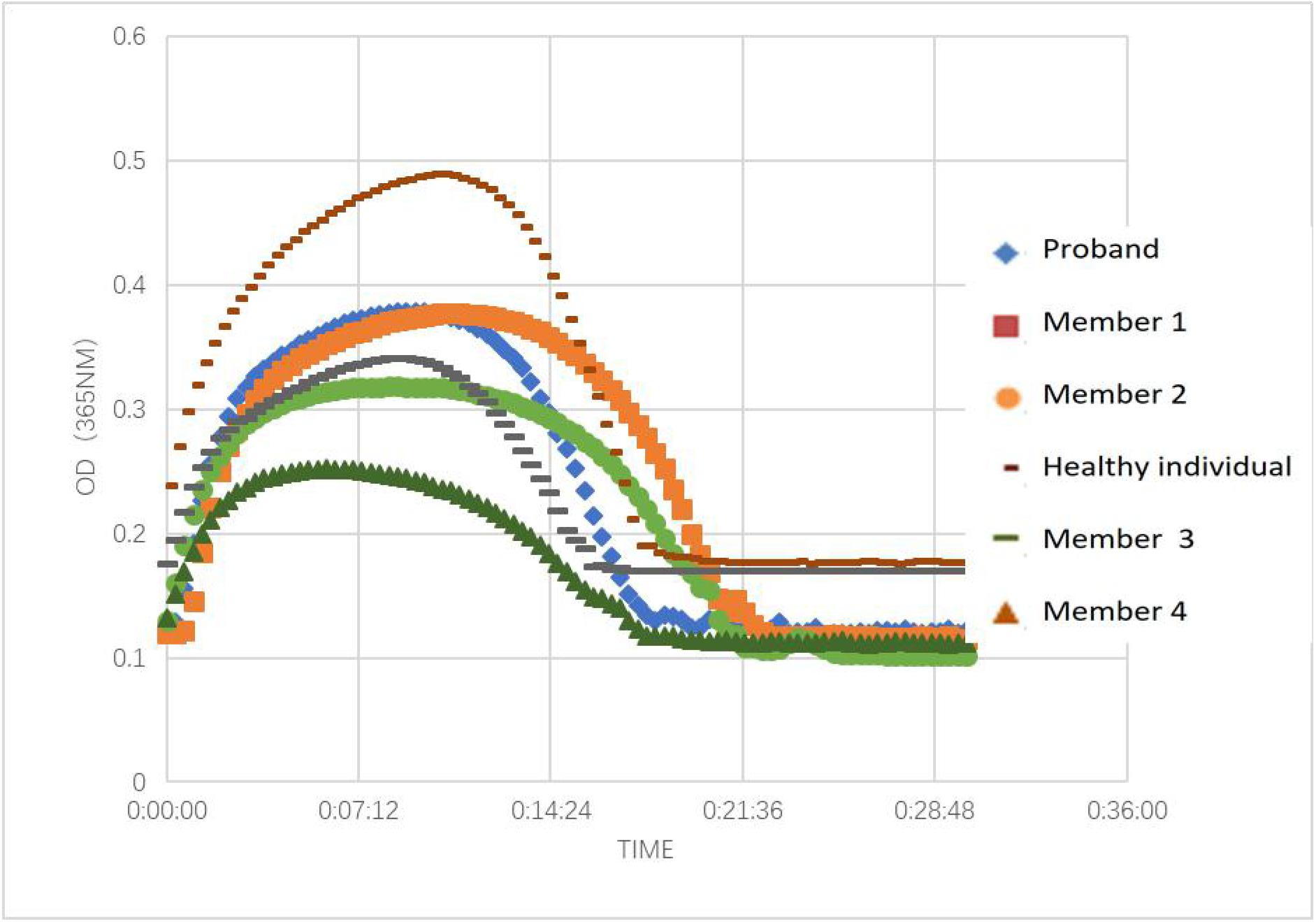
Fibrin clot lysis curves.

### Thromboelastography

The K values (time when the strength of blood clot reaches 20 mm) of patients with CD were increased; the average value was 3.7 min for the patients whereas it was 2.3 min for healthy individuals. Further, the Angle values (the angle between tangent line and horizontal line from the point of clot formation to the maximum curve radian of the figure) were decreased. The average value was 52.8° for the patients, whereas it was 61.5° for the healthy individuals. The thromboelastography results are shown in Figure 5 and Table 2.

**Figure 5.**
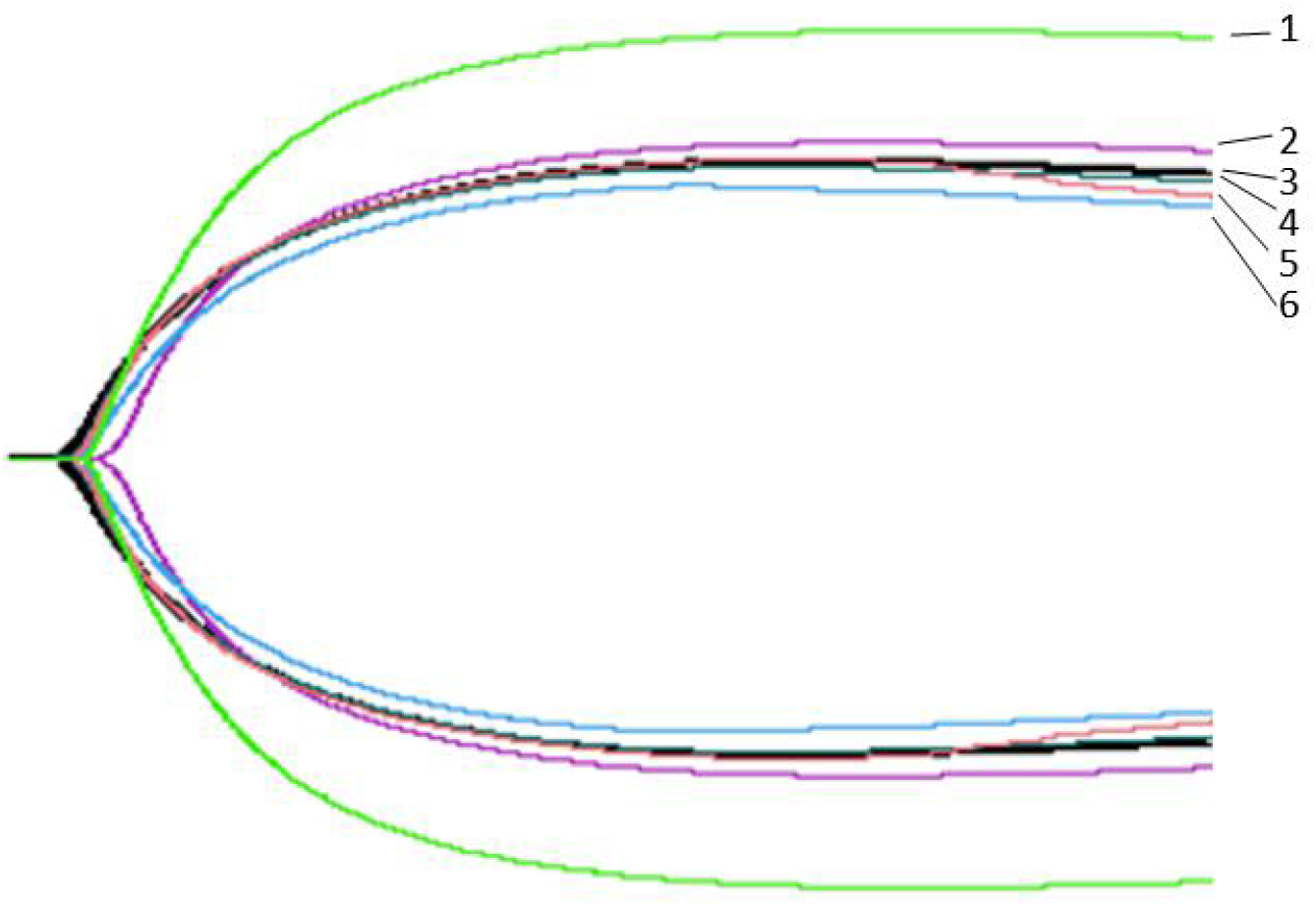
Thromboelastography. 1: Healthy individual; 2-6: Proband and her family members.

**Table 2.** Thromboelastography in the proband, her family members, and healthy individual

| Object | K (min) | Angle (°) | CI |
| --- | --- | --- | --- |
| Healthy individual | 2.3 | 61.5 | 0.9 |
| Proband | 4.1 | 53.1 | -2.0 |
| Member 1 | 3.2 | 52.7 | -2.6 |
| Member 2 | 3.4 | 53.2 | -1.8 |
| Member 3 | 3.5 | 54.3 | -1.8 |
| Member 4 | 4.3 | 50.9 | -3.0 |
| Reference Range | 1-3 | 53-72 | -3-3 |

Note: K value is the time when the strength of blood clot reaches 20 mm; Angle value is the angle between tangent line and horizontal line from the point of clot formation to the maximum curve radian of the figure, CI: comprehensive coagulation index

### SDS-PAGE

The SDS-PAGE results revealed that the proband and her family members did not show any variant protein bands (Figure 6).

**Figure 6.**
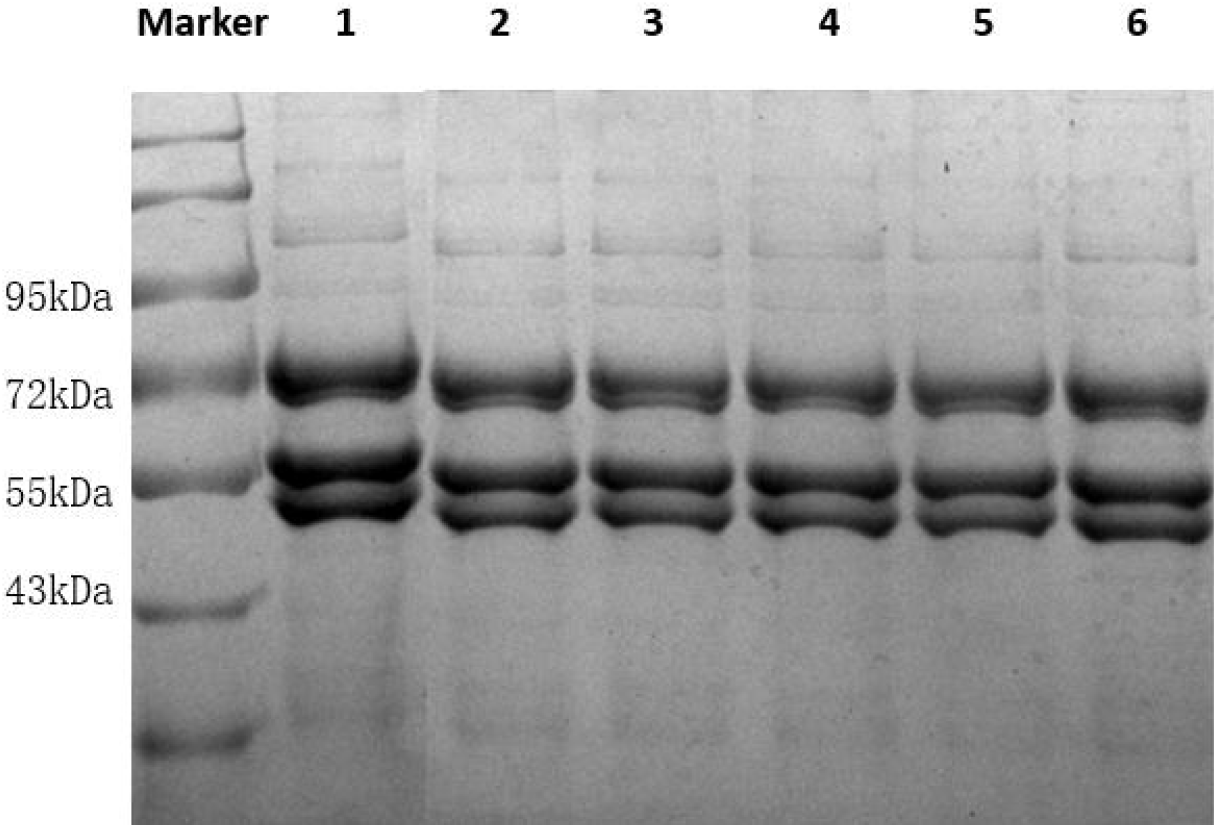
SDS-PAGE of fibrinogen. 1: Healthy individual; 2-6: Proband and her family members.

### Ultrastructure of fibrin clot

Shown in Figure 7 (A) is the structure of the control fibrin clot under a scanning electron microscope. The diameter of the fiber filament was uniform, the fiber nodes were few, the arrangement was neat and compact, the fiber filament interweaved and overlapped to form a fiber network, the spatial structure was relatively dense, and the network pore size was small. (B) The fibrin clot network structure of the proband showed that the fiber filaments were of different thickness, arranged irregularly, and the ends of the fiber filaments were curled into a mass. The fiber network space structure was loose, the network aperture increased, and the fiber branch nodes were more than those in the healthy individual.

**Figure 7.**
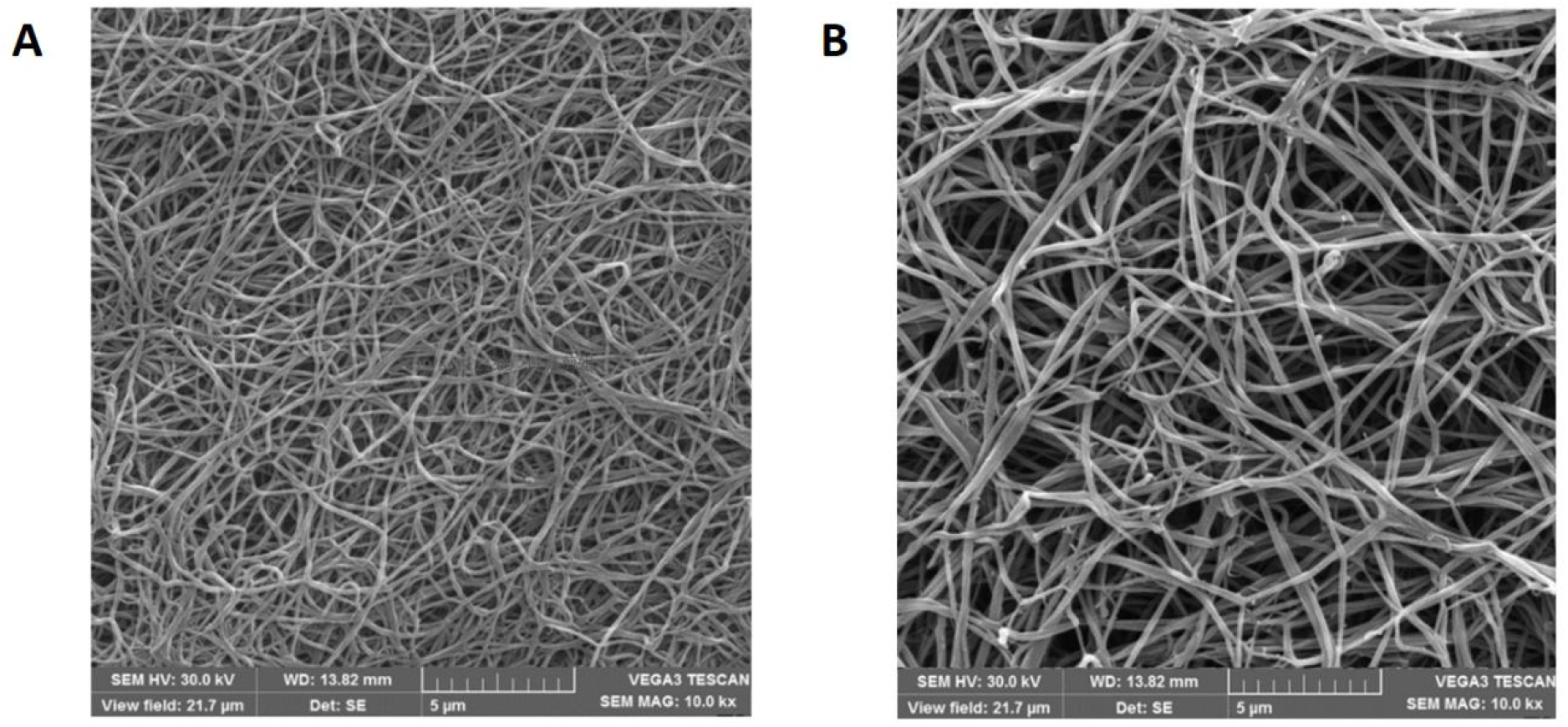
Scanning electron microscopy of fibrin clot. A: Healthy individual; B: Proband.

### Modeling and analysis of amino acid mutations

In the molecular model of wild-type fibrinogen, γ Ala327 is located in the α helix of D domain of the fibrinogen γ chain. The α helix is crucial to maintaining protein stability. The γ Ala327 backbone and γ Ser332 backbone form a hydrogen bond, which is 3.19 nm long. When Ala is replaced by Val, the hydrogen bond between γ-Val327 and γ-Ser332 did not change, but the side chain became longer, which affected its spatial structure, and the electrostatic force around it changed, which led to a change in the structure of the α-helix and weakening of protein stability (Figure 8).

**Figure 8.**
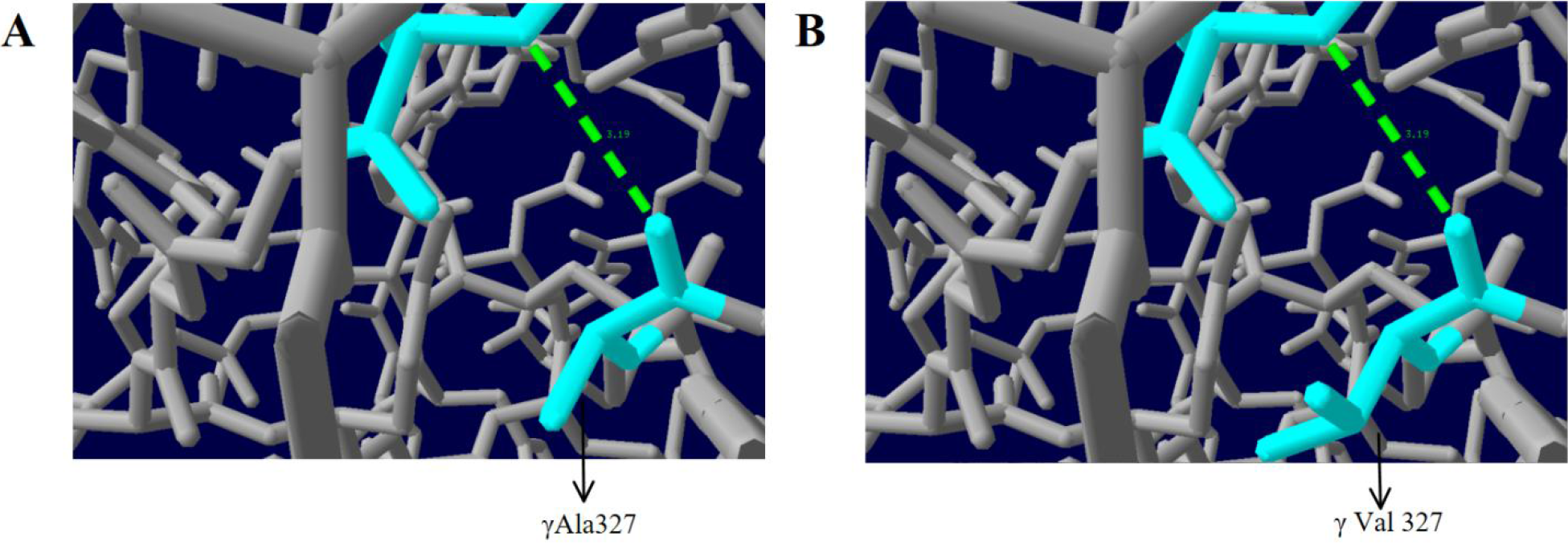
Analysis of the γAla327Val mutation with protein modelling. A: Healthy individual; B: Proband.

## Discussion

Congenital dysfibrinogenemia (CD) is a hereditary blood disease caused by the defects in fibrinogen genes, which leads to the abnormal structure and function of fibrinogen, and may affect coagulation. Fibrinogen genes (FGA, FGB, and FGG) are located on chromosome 4 of the human genome, and mutations in any of them may cause changes in the structure and/or function of fibrinogen. The mutations in fibrinogen gene include missense mutation caused by single base substitution, frameshift mutation caused by single base insertion or deletion, and mutations in the regulatory region. Single base substitution is the most common form of mutation that results in CD. So far, over 450 mutations in fibrinogen gene has been registered (www.geht.org). Single amino acid mutations are reported at 212 sites, of which mutations in FGA gene are the most common followed by mutations in FGG [8]. AαArg16 and γArg275 are the most common mutation sites in CD, and most common substitutions are AαArg16His, AαArg16Cys, γArg275Cys or γArg275His, which can be used as gene sites to screen for CD. In 101 cases of CD reported, mutations at these two sites accounted for 74% of the mutations [9].

In this study, the genetic testing results of the proband and her family members revealed a heterozygous missense mutation (C. 1058C > T) in exon 8 of the FGG gene, which resulted in Ala to Val substitution at position 327 of the γ chain. Structure modeling analysis showed that this change resulted in a spatial position and electrostatic force change, resulting in change in the structure of the α-helix and weakening of stability. Although interaction between D:E regions is the main driving force of fibrinogen aggregation, the D:D interaction is also important for fibrin oligomer formation [10–12]. The role of D:D interaction is to guide the D regions of the two fibrinogen monomers to connect; D:D interaction thus assists γ chain polymerization, which is necessary for fibrin end-to-end binding. By projecting the D:D region and analyzing its crystal structure, it was found that the interface of D: D polymerization is composed of hydrogen bonds between two adjacent fibrinogen molecules, and that the D: D interaction is crucial for fibrin monomers to aggregate horizontally and linearly [10, 13]. The amino acids at position 88-406 of the gamma chain encoded by FGG gene form the main functional domain of the D region [14], and the mutation site γAla327 investigated in this study is located in this region. In γAla327Val, alanine is replaced by valine, which affects the secondary structure, changes spatial arrangement, affects the D:D binding between the gamma chains of fibrinogen, and hinders the formation of fibrils, thereby affecting the aggregation of fibrinogen. Our aggregation test and thromboelastography results revealed that the proband and her family members had impaired fibrin aggregation.

The ultrastructure of the probands’ fibrin clots was observed by scanning electron microscopy. The sizes of the filaments were different, the arrangement was irregular, the ends of the filaments were coiled, the fiber network was loose, the aperture of the network was enlarged, and there were many fiber nodes. Defects in the D:D interaction between adjacent fibrinogen molecules leads to easy lateral extension of the filament, resulting in an increase in branch points and formation of thinner filaments [15]. Substitution of γAla327 results in incorrect end-to-end location of fibrin monomers during aggregation, resulting in the increase in fiber branching. Further, during polymerization, normal fibrin monomers can combine with mutant monomers, resulting in inconsistent fiber diameter, irregular arrangement, looseness, and large mesh-like structure. Analysis of the mutant fibrin clot network showed that the fibrin clots with impaired D:D interaction (γArg275His, γArg275Cys and γArg375Gly) had more pores in the network structure, and that the network comprised many conical fibers and pores [16]. These results are similar to those of this study. Sugo et al [14] suggested that the structure of fibrin clot might be related to the clinical phenotype, and the phenotype could be predicted by studying the ultrastructure and network structure of fibrin clot. Studies have shown that the dissolution rate of fibrin clot depends on the density of its structure rather than the diameter of fibrin fibers, and the loose fibrin network structure composed of crude fibers is easier to be decomposed than the dense fibrin network structure composed of fine fibers [17]. Patients with high fibrin density in the fibrin network have a higher risk of thrombotic events [18].

In patients with thrombotic CD, such as fibrinogen Paris V [19] (FGAc.1717C > T) and fibrinogen Perth [20] (FGAc.1541delC), fibrinolytic time is prolonged. In this study, fibrinolysis prolongation and fibrinolysis resistance were not found in patients with CD when compared with healthy individuals, suggesting that γAla327Val mutation does not affect fibrin clot dissolution.

The concentration of fibrinogen antigen detected by immunoturbidimetry was normal in patients with CD, but the concentration of fibrinogen activity detected by Clauss method was significantly decreased. Immunoturbidimetry is based on the specific binding of fibrinogen antibody to fibrinogen. As fibrinogen antigen determinants are present in mutant fibrinogen molecules, the concentration of fibrinogen antigen in plasma of patients with CD will not be reduced. Combination of multiple methods to detect fibrinogen can reduce the rate of misdiagnosis and missed diagnosis of CD. Our previous study showed that the combination of PT-derived method and Clauss method is helpful to screen CD [21]. In this study, in the proband, the concentration of fibrinogen activity was significantly lower than that of fibrinogen antigen (0.75 g/L vs 1.59 g/L, reference range: 2-4 g/L). Liver and kidney function, electrolyte, and blood routine examinations were not abnormal, excluding secondary factors. Gene sequencing found γ Ala327Val mutation, which could be used to diagnose the proband as having CD. Through fibrinogen aggregation test, fibrin clot dissolution test, and thromboelastography, the function of fibrinogen was analyzed, and the defect of fibrinogen aggregation function was found. The ultrastructure of fibrin network was observed by electron microscopy, and the changes in fibrinogen structure caused by the mutation were studied. It was found that the branches of fibrin network increased, and the arrangement was irregular, loose and the mesh was large. These changes may affect the process of fibrin aggregation. Although no bleeding or thrombotic events have occurred in this family, the possibility that thrombotic or hemorrhagic events will occur in the future, especially in the case of surgery, postpartum, or trauma, cannot be excluded. Long-term follow-up studies of patients with CD have shown that patients without clinical symptoms at diagnosis may still have thrombotic or bleeding events several years later, and that their risk of bleeding or thrombosis is relatively high [9]. Barbara et al [22] reported a case of CD with an α Arg16His homozygous mutation (fibrinogen Giessen I), who had no bleeding or thrombotic events in daily life, but had symptoms of massive bleeding during delivery. Therefore, the proband and her family still need follow-up observation to prevent thrombosis or bleeding and other events.

## Acknowledgments

We would like to thank the Department of Hematology and Clinical Laboratory of the First Affiliated Hospital of Guangxi Medical University, China. This research was supported by Grants from the National Natural Science Foundation of China (81560342).

## Author contributions

The authors declare that they have no conflict of interest with the content of this article.

